# Horizontal transfer in bacterial Methionyl tRNA synthetase is very common shown by Genus and phyla level phylogenetic analysis

**DOI:** 10.1101/042366

**Authors:** Prabhakar B. Ghorpade, Avinash D. Pange, Bhaskar Sharma

## Abstract

Methionyl tRNA synthetase is single copy informational gene in Salmonella typhimurium. Informational genes are more conserved than operational genes. In this study we had analyzed HGT events within MetG sequences of different bacterial genera. A species tree based on 16srRNA sequences of the same genus was drawn evaluated against the generally accepted species tree of the bacteria. MetG phylogenetic tree was evaluated against the 16srRNAS tree and HGT event identified. Similarly phyla trees were made and HGT event identified. 24 HGT events were identified between genus and 11 within phyla. MetG is a considered as conserved gene finding so many HGT event in this gene indicate that horizontal gene transfer is very common in this gene. Manual tree making for phyla could help to understand phylogenetic relationships between very large trees.

## Background

Phylogenetic trees constructed from different genes for same set of organism often gives different phylogeny. The difference in phylogeny could be due to insertion, deletion, sampling errors, differences in rate of substitution, lateral gene transfer etc within these genes; making it difficult to choose a gene to reconstruct phylogeny[1]. In higher animals mitochondrial genes are often chosen because these genes are inherited together and no recombination events are known between them [2]. In case of microorganism ribosomal genes are the key standard for phylogenetic analysis because 16srRNA genes undergo least HGT between species and are of sufficient length (1500 bases) contains both conserved and variable regions to enable species comparison[3-5]. Authentic species trees are used for tracing phylogeny[6] and also are for identifying horizontal gene transfer (HGT) between species [4]and phyla.

Horizontal gene transfer can be identified based on different codon usage pattern or GC content for a particular gene in comparison to other genes in organism [7]. Other approach involves comparison of gene specific tree with species or real time tree. In comparing the 16SrRNA is considered standard for that organism and treated as species tree then one can compare gene of interest phylogeny as gene tree[8, 9]. HGT is lateral gene transfer between different species[10, 11]. The HGT between different species can be explained at phylum level by treating any bacterial species showing up in displaced location as separate phylum.

In bacteria horizontal transfer is a common phenomenon affecting true phylogeny[12], despite these problems species trees for bacteria have been drawn using multiple conserved gene. This tree is generally accepted as representing true phylogeny within bacterial species [4, 6,13].

Tree topologies generated by methods using different statistical computation are different. Generally trees are character based or distance based. Character types include Maximum Parsimony, Maximum Likelihood and Bayesian inference and distance methods include Minimum Evolution and Least squares. The clustering algorithm with distance as data type, have UPGMA, Neighbourhood Joining and Fitch-Margoliash as methods to draw phylogeny[14-17]. Studies comparing the trees generated by different methods with real time tree shows different fit with the true species tree.

Methods to compare different phylogenetic trees are based on dissimilarity or common information measurements. The dissimilarity methods include Robinson-Foulds (RF)[18], Tree-Bisection-Reconnect (TBR), Subtree-Prune-Regraft (SPR), Geodesic Tree Distance[19], Tanglegrams[20] and Rotation Distance [21].The Common information model for tree comparison includes Maximum Common Refinement Subtree and Maximum agreement subtree[22].

In bacteria generally accepted species tree is that of16srRNA which is largely based on ribosomal RNA sequences [5]. Ribosomal RNA sequences are conserved but in recent studies it has been shown that they too are subjected to HGT. Another conserved gene is methionyl tRNA synthetase, only single copy of this gene is present in Salmonella enetica serovar typhimurium. Aminoacyl tRNA synthetases (AARS) are important for supplying amino acid charged tRNAs to facilitate translation. AARS are classified into Class I and ClassII based on differences in their catalytic sites [23]. MetG is classified into Class I owing to the presence of HIGH and KMSKS signature as amino acid sequence[24, 25]. Among AARS Methionyl tRNA synthetase (MetG) is required for initiator tRNA and supplying Methionine charged tRNA.

In this study we have taken 57 Bacterial Genus representing all the phyla of bacteria mentioned in Hugenholtz phylogeny [26, 27]and downloaded the 16srRNA [28]and MetG sequences (NCBI). Species tree was drawn based on 16srRNA sequences and this species tree was compared with accepted species tree[26]. The tree which gave minimum RF distance was taken as species tree and was used as species tree for finding HGT within species in MetG generated tree. The results are discussed specifically in the context of MetG and in general.

## Results

### Sequences

Sequences and accession no of sequences of MetG and 16srRNA used in this study are given in supporting file1 and file 2.

### Trees created

The 16srRNA and MetG trees created by ML,MP, ME, NJ, UPGMA are available in supporting files(File 3).

### Species tree

The topological distance between tree generated by different methods and Hugenholtz phylogenetic tree (Fig1) as realtime tree are shown in Table1. Minimum Evolution (ME) and NJ tree gave lowest value of RF distance. These trees were identical trees since RF distance between them was zero. 16srRNA ME tree was taken as species tree for further analysis.

**Table 1.** Topological distance between Hugenholz tree and 16srRNA Trees

| **Trees** | **Robinson Fould distance** | **Least-squares coefficient(LS)** | **Bipartition dissimilarity** |
| --- | --- | --- | --- |
| ML16srRNA | 59 | 287613602416.595 | 93.5 |
| MP16srRNA | 58 | 1900590657932.631 | 109.5 |
| **ME16srRNA** | **55** | **287613563786.531** | **88.0** |
| NJ16srRNA | 55 | 287613563786.531 | 88.0 |
| UPGMA16srRNA | 59 | 287613567458.186 | 93.0 |

**Fig.1.**
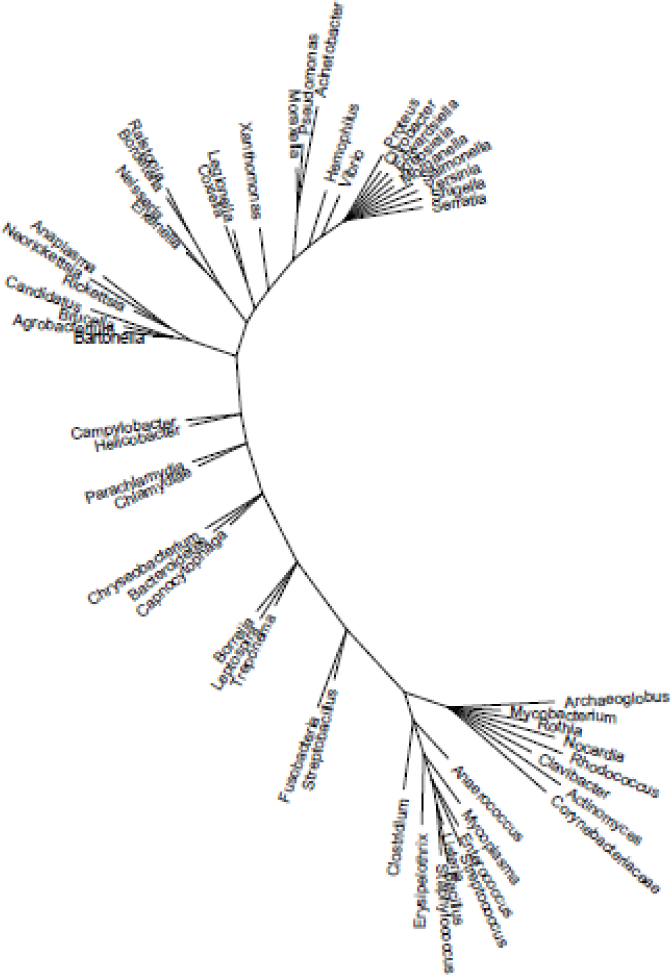
Hugenholtz tree representation by Tree builder

### MetG tree

Topological distance between various trees generated with MetG sequences and the species tree 16srRNA ME selected above are shown in table 2. The minimum topological distance with of all the methods was found between Maximum parsimony tree of metG and the species tree of 16srRNA.

**Table2a.**
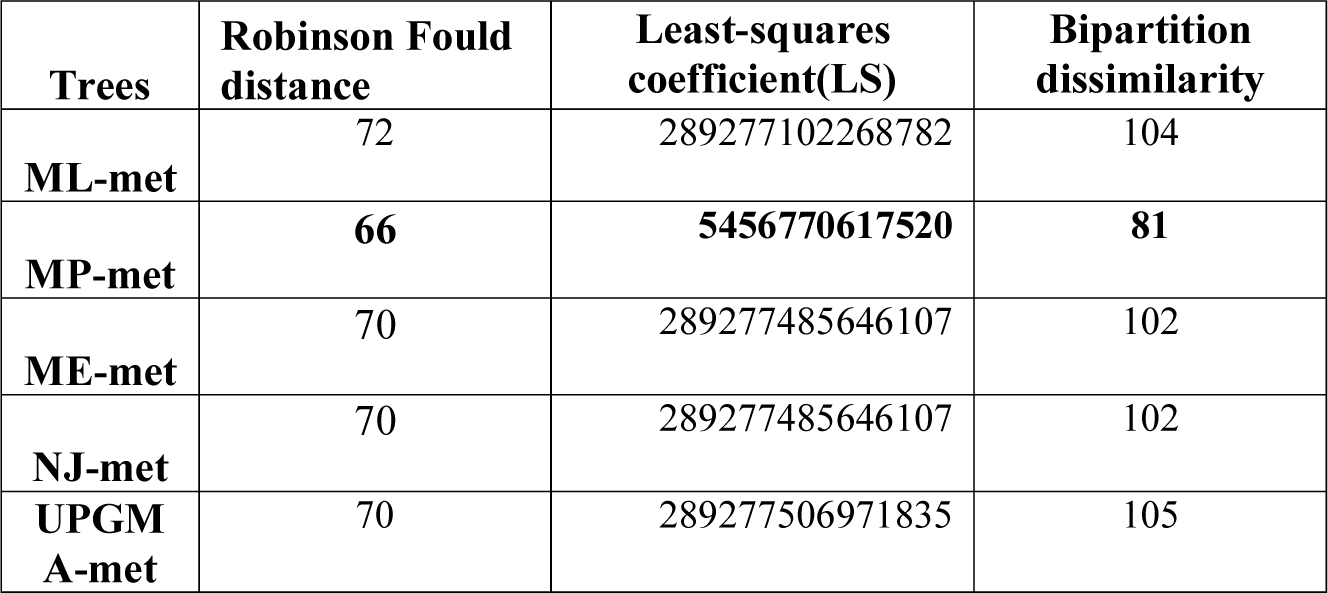
Topological distance between 16srRNA tree and metGtre

### Phyla tree

Phyla trees were generated by choosing the option collapse in Figtree and drawing phyla in tree builder. Phylum tree created in tree builder and viewed with tree viewer of T-REX for 16SrRNA is shown in Fig.4; for metG phylum tree is shown in Fig.5.

### HGT

The HGT was calculated between phyla and species trees. As was expected the HGT within phyla tree was less 11 HGT (Fig.7) than species tree 24 HGT (Fig.6).

## Discussion

With the increase in the number of bacterial sequences, our understanding of evolutionary relationship between bacterial genera tends to change. The new sequences when included for generating a phylogenetic tree may cause addition of clades in established or new location this may change relevance or irrelevance of phylogeny within established trees; and thus give new understanding to relatedness within genus, phyla or kingdom or even universal tree.

Generally accepted bacterial phylogeny is that of Hugenholtz [26, 27]. We had used that as species tree to evaluate 16srRNA tree generated in this study assuming that it would represent true species tree and by comparison with that we can find HGT in metG sequences. 16sRNA sequences are generally accepted as conserved with no HGT but recently it has been shown that these sequences are not as conserved as assumed and HGT within 16srRNA sequences is present[35]. We used distance as measurement of dissimilarity between various trees.ME and NJ 16srRNA trees gave minimum dissimilarity with the species tree. ME [36]and NJ [37]generally give identical tree if the number of sequences are less [38-40]. A comparison of real time sequence tree withdifferentphylogenetic trees for HIV reveal that best fit tree by phylogenetic methods were NJ, ML [40]. Based on these results we took ME 16srRNA tree as species tree for HGT estimation in MetGsequences.Comparison of distance measurement with the trees generated by different methods for MetG sequences was used to select the best fit tree. This time tree generated by Maximum parsimony gave lowest RF distance. Maximum parsimony tree is generally used for protein sequences and since protein sequences was used for generating MetG tree that could be the reason for getting lowest RF distance between MetG and 16srRNA species tree generated by us. It is difficult to predict true species tree from the different tree generating methods[40]. Alignment with a known species tree is the true measure for making such prediction.

We found 24 HGT events within MetG sequences, which is on the higher side considering that this protein is a product of informational gene and has high modular structure. The reasons for higher HGT event we observed are difficult to explain. One reason which may partly explain this anomaly could be the species selection. The number of clades/branches in the tree generated by us with MetG sequences had shown different branches particularly proteobacteria (Fig. 3, 5) whereas these were together in Hugenholtz tree (Fig1.) and in 16srRNA tree (Fig.2, Fig4). The high incidence of HGT in MetG sequences found in this study suggests that a relook on prevalence of HGT in other conserved gene sequences is needed.

**Fig.2.**
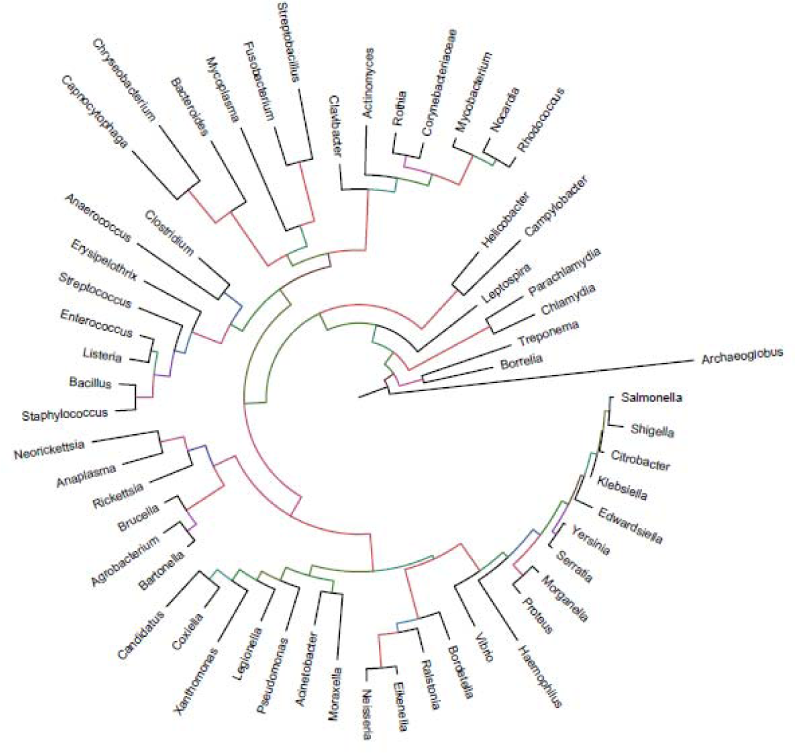
The 16srRNA Minimum Evolution tree

**Fig.3.**
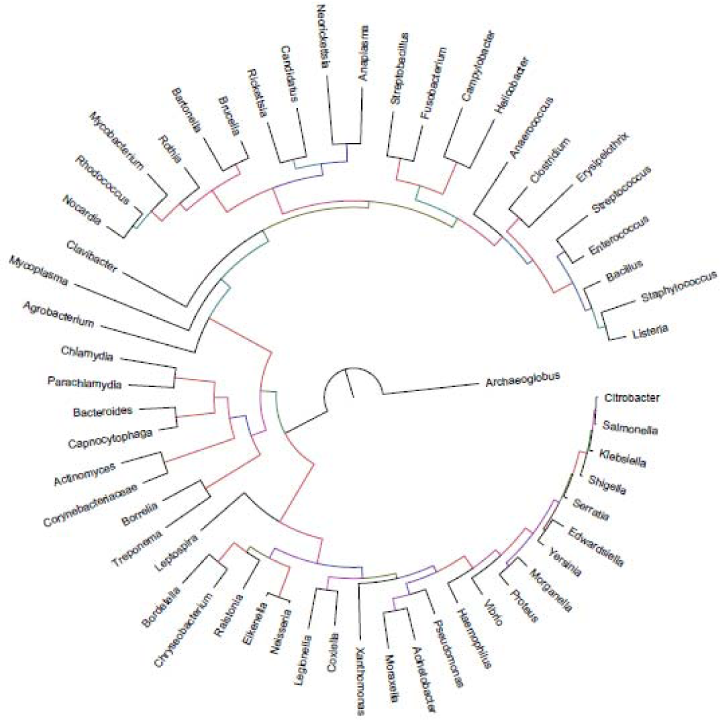
The metG tree made by Maximum Parsimony method

**Fig.4.**
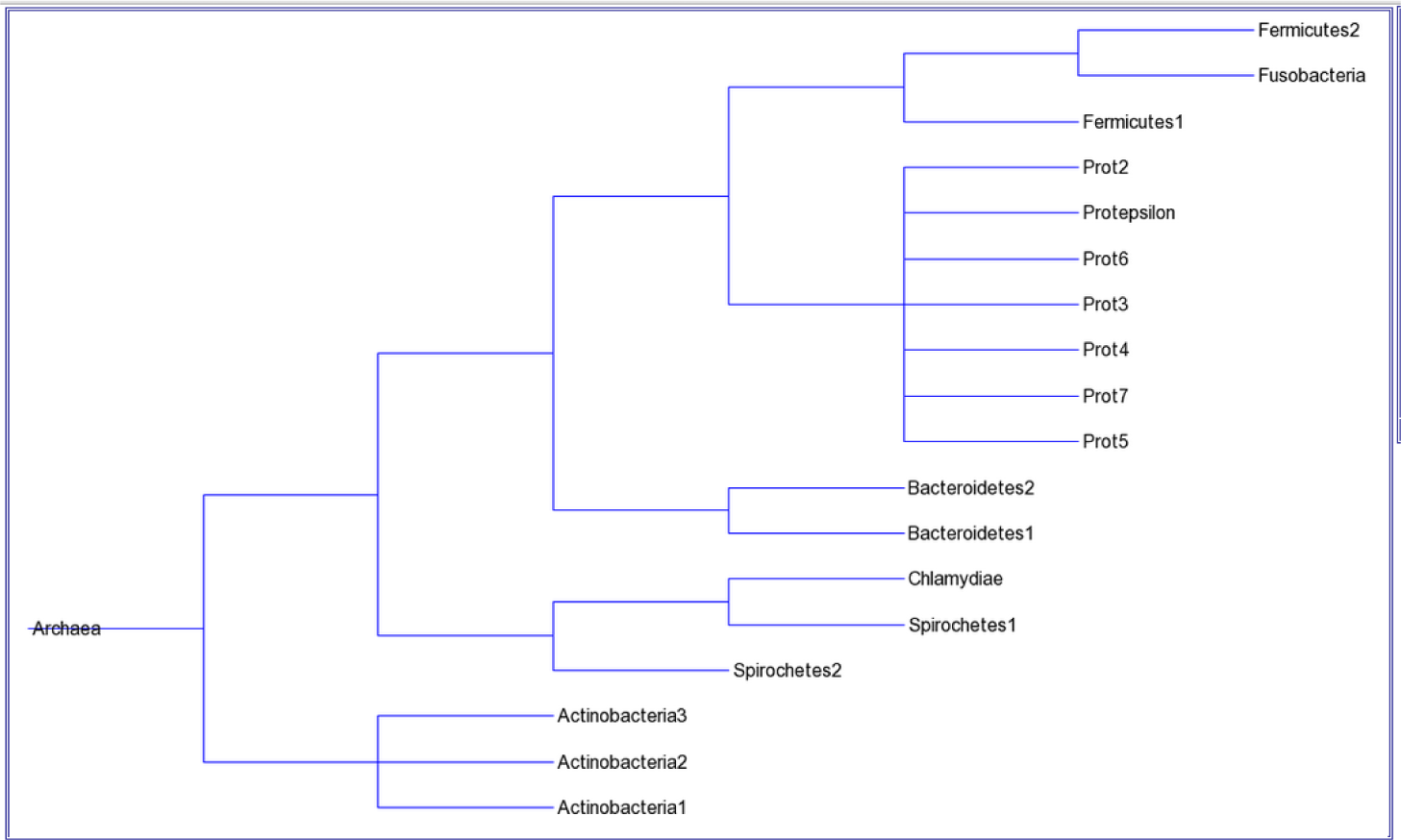
16srrna Phylum tree made in Trex tree builder

**Fig.5.**
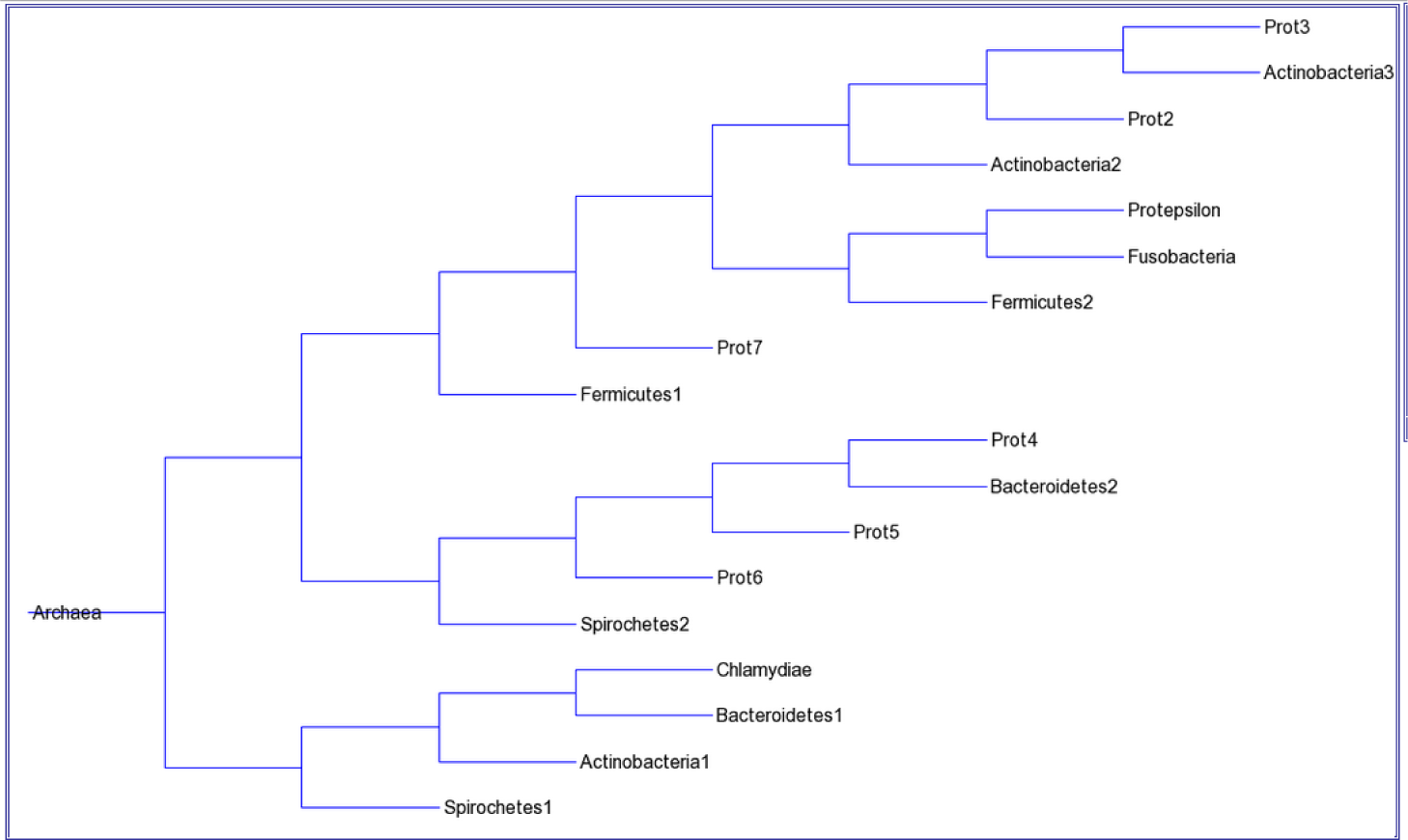
metG(MP) Phylum tree made in Trex tree builder

**Fig.6.**
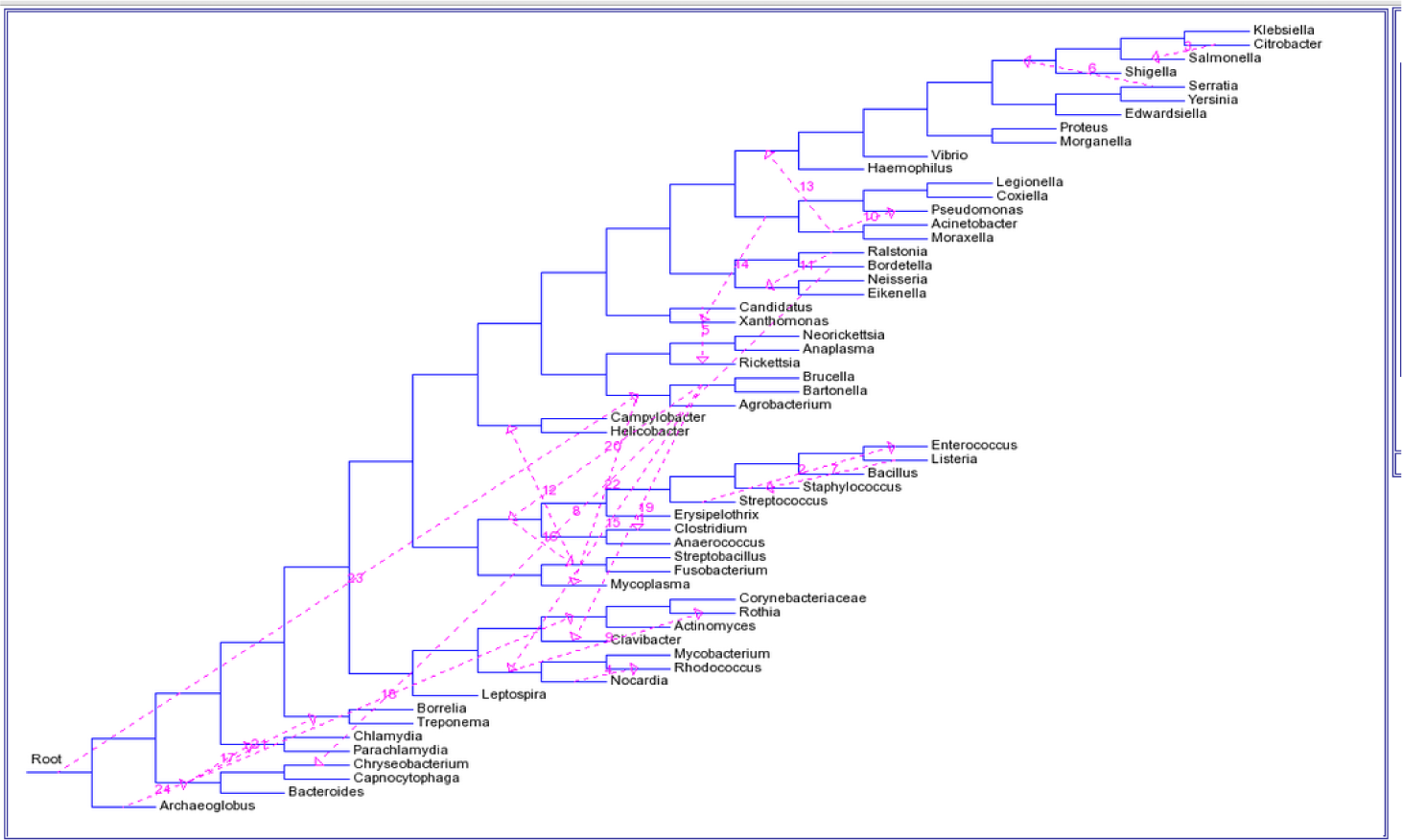
Horizontal transfer in genus of 16SrRNA tree made by Minimum Evolution (species tree) with metG tree made by Maximum Parsimony method (gene tree)(HGT=24)

When the HGT was compared between phyla 11 HGT (Fig.7) incidences were found, this suggest that HGT within phyla is less common than genus. It has been suggested that HGT is dependent on evolutionary distance [12, 41-44].

**Fig.7.**
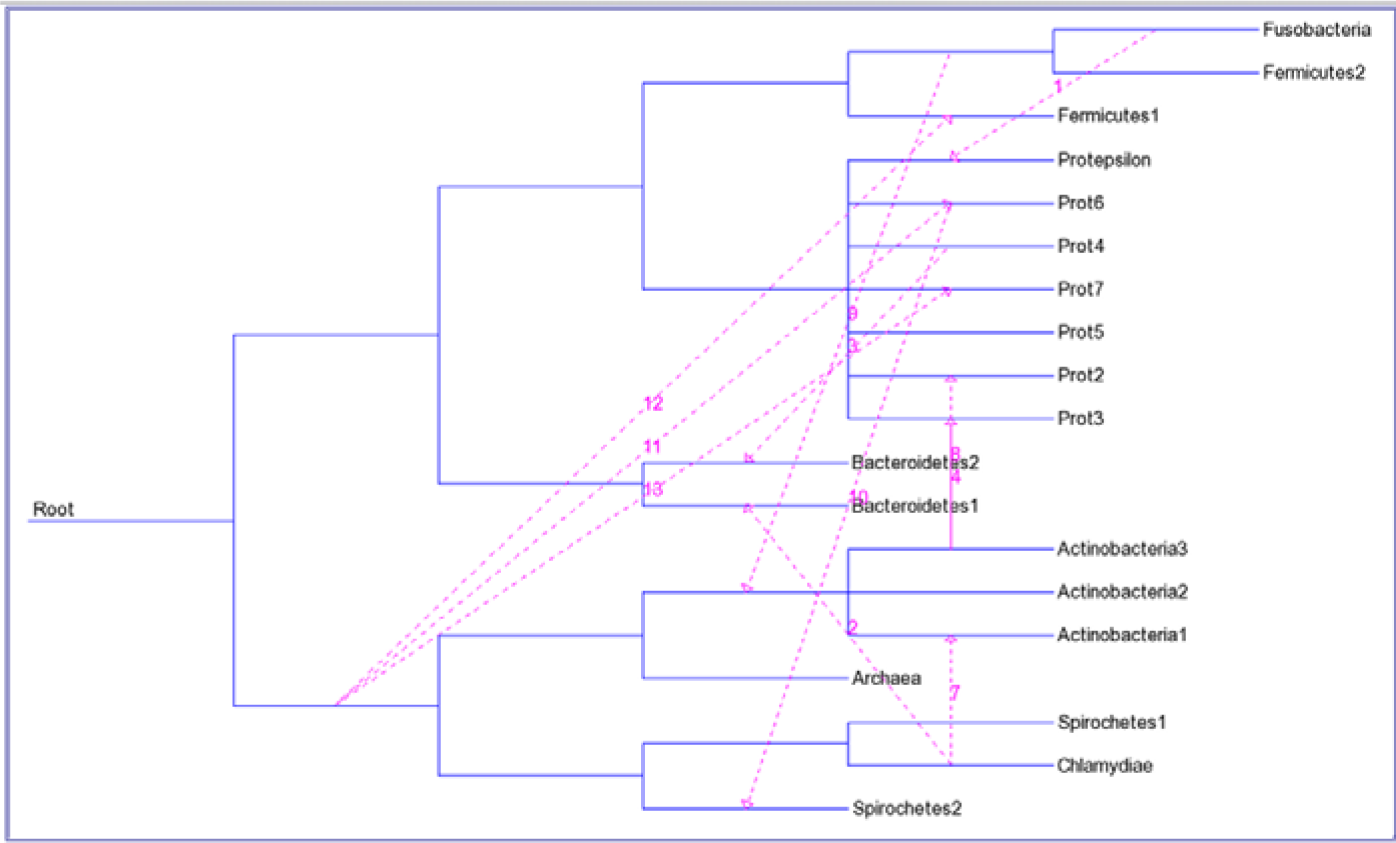
Horizontal transfer in genus of 16SrRNA Phyla tree (ME) (as species tree) with metGPhyla tree (MP) (gene tree)(HGT=11)

Our results are in accordance with this fact. This is the first description of HGT within phyla and genus tree. Phyla level HGT by virtue of its relative simplistic network in comparison to genus network may help to understand jumbled network created by intensive gene transfer events which some author predict could replace classical rRNA based phylogeny[10, 11, 45, 46].

## Materials and Methods

### Retrieval of MetG Sequences

Salmonella enterica subsp. Entericaserovar Typhimurium genome sequence and other 57 amino acid sequences in fasta file format were retrieved from NCBI.

The 16SrRNA for creating species phylogeny for corresponding genus mentioned in metG sequences were downloaded from Ribosomal Database-Silva from www.arb-silva.de [28].

### Alignment of MetG and 16SrRNA sequences

The alignments of sequences were performed in MEGA [29] using Muscle. Unaligned region of alignment were removed.

### The Evolutionary analysis

Evolutionary analysis was done in MEGA [30-32] and the methods used were Maximum Likelihood tree, Neighbour Joning tree, Minimum Evolution tree, Maximum Parsimony tree and UPGMA.

Test of reliability was Bootstrap method[33]with 500 replications. In order to keep things simple we have kept captions givenin MEGA with minor changes for each figures to get information regarding methodology of tree analysis.

### Species tree

The framework of Hugenholtz species tree as given in [26, 27], was used to construct corresponding species tree in Tree builder canvas of T-REX with addition of archaea as root.This tree was used to select 16srRNA species tree for identifying HGT events.

### Analysis of Horizontal Gene transfer

The T-REX [34] was used for comparing tree generated by different methods. In the first step the tree corresponding to Hugenholz tree generated above was treated as standard species tree against 16srRNA gene tree generated by different methods. Then the 16srRNA gene tree showing lowest Robinson Foulddistance was selected as species tree foridentifying HGT events in MetG tree. MetGtree was drawn by different methods and each tree was compared with 16srRNA species tree. The tree which had shown smallest RF, least square and bipartite distance was used for identifying HGT within MetG sequences.

### Collapsing species into Phylum

The bacterial species belonging to same phylum and falling on same cluster were collapsed into phylum in Figtree(http://tree.bio.ed.ac.uk/software/figtree). This action was performed on ME tree of 16SrRNA and MP tree of metG sequences.

### Phylogenetic tree at Phylum level

Figtree was used to get collapsed phylum tree with individual species on branch represented as phylum by annotating. Then T-REX Newik Builder 0.3 was used to build the phylogenetic tree for phylum level representing corresponding species phylogeny.

### Viewing Phylogenetic tree

Manually drawn phylogenetic tree was viewed with Tree Viewer from T-REX server.

### Analysis of Horizontal Gene transfer at phylum level

HGT was calculated between species tree (16SrRNA ME tree) and gene tree (Met MP tree) in Trex server.

## Acknowledgement

We thank Indian Veterinary Research Institute for supporting this work. We also thank ICAR (GP, PA, BS) and CSIR (GP) for financial support for this research.

